# Three Rules Explain Transgenerational Small RNA Inheritance in *C. elegans*

**DOI:** 10.1101/2020.01.08.899203

**Authors:** Leah Houri-Ze’evi, Olga Antonova, Oded Rechavi

## Abstract

Life experiences trigger transgenerational small RNA-based responses in *C. elegans* nematodes. Dedicated machinery ensures that heritable effects would re-set, typically after a few generations. Here we show that isogenic individuals differ dramatically in the persistence of transgenerational responses. By examining lineages composed of >20,000 worms we reveal 3 inheritance rules: (1) Once a response is initiated, each isogenic mother stochastically assumes an “inheritance state”, establishing a commitment that determines the fate of the inheritance. (2) The response that each mother transfers is uniform in each generation of her descendants. (3) The likelihood that an RNAi response would transmit to the progeny increases the more generations the response lasts, according to a “hot hand” principle. Mechanistically, the different parental “inheritance states” correspond to global changes in the expression levels of endogenous small RNAs, immune response genes, and targets of the conserved transcription factor HSF-1. We show that these rules predict the descendants’ developmental rate and resistance to stress.

---

Even when environmental conditions are tightly controlled, populations of isogenic *Caenorhabditis elegans* nematodes exhibit inter-individual differences in many traits including developmental timing, response to stress, behavior, and life span [1–7]. Further, worms display substantial variability in the manifestation of heritable epigenetic effects [8–10].

*C.elegans* can control the expression of genes transgenerationally via heritable small RNAs [11–15]. RNA interference (RNAi) and RNAi inheritance are modulated by different external challenges [6,16–21], and even by internal neuronal processes [22]. Heritable small RNAs are amplified in the next generations by specialized RNA-dependent RNA polymerases and thus avoid “dilution” [17,23].

Exogenous introduction of double stranded RNA (dsRNA) triggers transgenerational silencing of germline genes [9]. Typically, heritable RNAi is maintained for ~3-5 generations (the so-called “bottleneck” to RNAi inheritance [9]), unless it is re-set earlier by stress [24]. While this is the usual duration of heritable responses at the population level, some individuals continue to silence genes for multiple generations, and others rapidly lose the inherited effects. It has been shown that selection enables maintenance of heritable RNAi indefinitely (>80 generations) [10]. Control over the duration of transgenerational silencing is achieved via an elaborate regulatory pathway [8], and disabling some of the genes in the pathway (for example, *met-2,* which encodes for an H3K9 methyltransferase) leads to stable inheritance [14,25,26]. Currently, the source of variability in the inheritance capacity of genetically identical individuals is completely unaccounted for.

To examine how heritable silencing is segregated across a lineage and distributed in the population, we exposed worms containing a germline-expressed single-copy *gfp* transgene to anti-*gfp* dsRNA. We scored *gfp* silencing in sister worms that derive from a single mother that was exposed to RNAi (we tightly controlled the exposure to RNAi and allowed the mothers to lay eggs for 6-8 hours, see **Methods**). As expected, 100% of the worms that were directly exposed to RNAi (the P0 generation) silenced the *gfp*. We then examined how these isogenic sisters pass on the silencing response transgenerationally. For this purpose, we used two different experimental setups: In the first set of experiments we tracked silencing across multiple generations, by propagating a few randomly selected worms from each lineage in every generation (**Fig. 1A** and **Methods**). In the second set of experiments, we monitored heritable silencing for one generation only, and scored the progeny of the entire synchronized RNAi-exposed population (**Fig. 1B** and **Methods**).

**Fig. 1.**
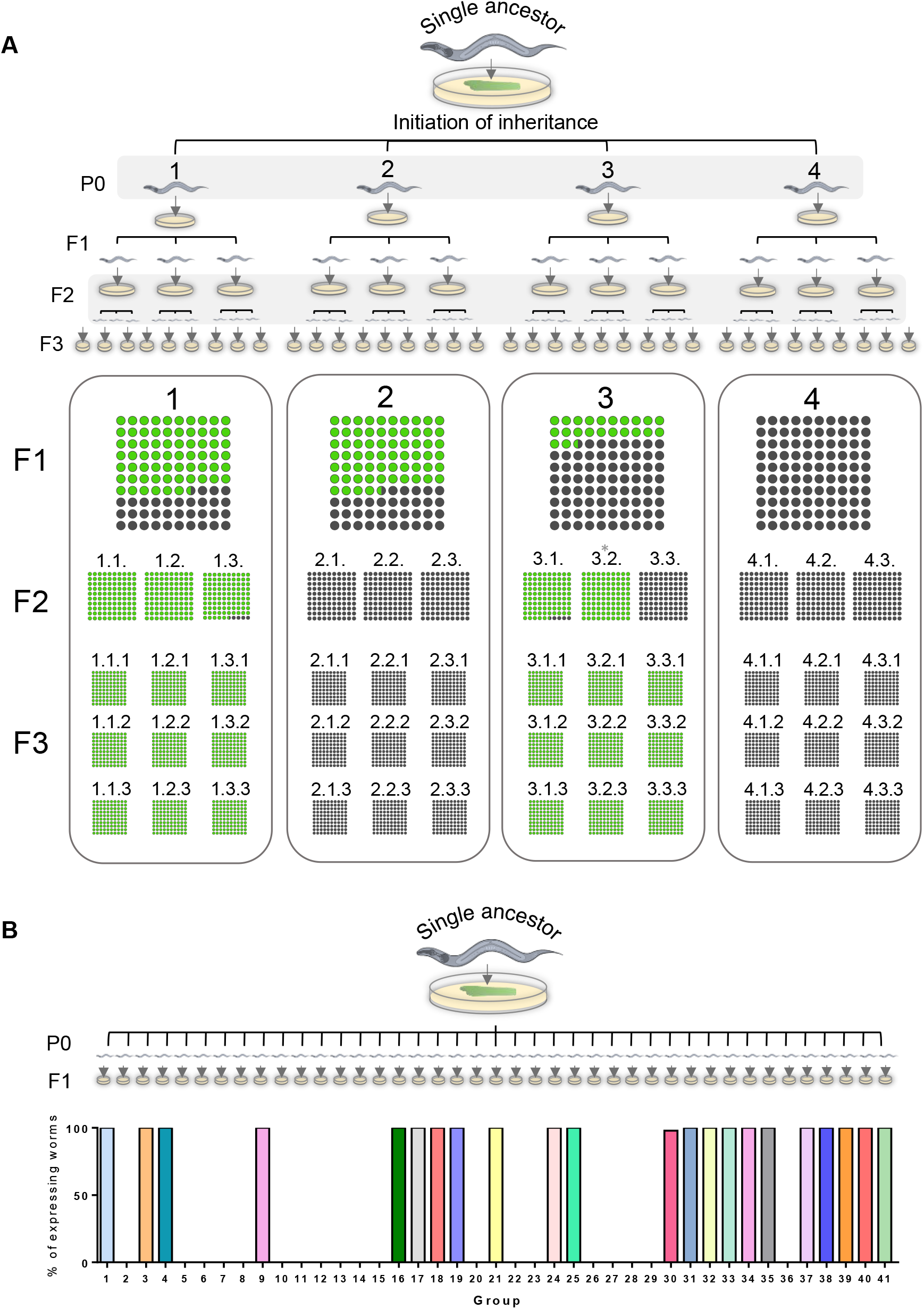
Inheritance patterns of an *anti-gfp* silencing response reveal striking uniformity in heritable silencing in worms which are derived from the same mother. A. Four P0 mothers, derived from one single ancestor, initiate an anti-*gfp* RNAi response. At each generation, the inheritance phenotype of the entire synchronized progeny of each mother is scored, and three worms are randomly selected to generate the next generation. Displayed are the proportion of worms that still inherit anti-*gfp* silencing response (dark circles) and worms that lost the silencing (green circles) in each group of progeny. The number of worms in each group of progeny ranged from 25 to 90. Mean = 43, median = 41.5. (*In group 3.2. only 7 worms were examined due to partial sterility of the mother.) B. Multiple (41) synchronized P0 mothers, derived from one single ancestor, initiate an anti-*gfp* RNAi response. In these sets of experiments the *in-utero* exposure of the F1 generation to the dsRNA trigger was further minimized. The inheritance phenotype of the F1 progeny of each mother is scored and presented as the percentage of worms that lost heritable silencing response (re-express GFP) in each group. Each color represents a separate P0-derived progeny group. For further details regarding synchronization and exposure times, please refer to the **Methods** section.

To our surprise, in the first set of experiments we found that in every generation individuals descending from a given lineage (from a single RNAi-treated mother), exhibit striking uniformity in heritable silencing levels, which became complete in later generations (**Fig. 1A**), regardless of the day in which the progeny were laid (namely it did not depend on the mother’s age). In the second set of experiments, we further controlled differences in *in-utero* exposure of the F1 progeny to the external dsRNA trigger (**Methods**) and thus completely eliminated heritable variability in the F1 generation. Still, when comparing multiple groups of progenies that were derived from different RNAi-treated P0 mothers, we observed strong differences in the dynamics and penetrance of the heritable silencing between the different lineages (**Fig. 1B**). In the past, RNAi experiments were traditionally performed by exposing populations of worms to RNAi followed by random sampling of worms in every generation[8,24,25]. Our experiments, in which we examined inheritance across lineages that originate in single mothers, reveal that the variability in the inheritance dynamics that is observed in population-based assays can be attributed to differences that begin in the RNAi-exposed mothers.

Why do isogenic mothers initiate different heritable RNAi responses? We reasoned that this inter-individual variability arises either because each mother is exposed to different amounts of dsRNA, or, more interestingly, because the RNAi inheritance machinery in isogenic mothers can assume different internal “states”. Despite extensive efforts to minimize potential differences in the exposure of individual worms to the dsRNA trigger (**Methods**), we still observed remarkable variability in the inheritance between lineages that derive from single isogenic P0 mothers. We thus examined whether this variability could instead be explained by internal interindividual differences in the activation state of each worm’s RNAi machinery. To this end, we analyzed the silencing patterns of a germline-expressed *mcherry* transgene, which gets spontaneously silenced, independently of exogenous dsRNA, by endogenous small RNAs [27]. Similar to the variability observed in dsRNA-induced anti-*gfp* silencing, we found that isogenic sister worms vary in their tendency to silence the *mcherry* transgene. Silencing accumulates over generations in the population, eventually resulting in a population of worms which all silence the transgene (see lineages of *mcherry*-silencing inheritance in **Fig. S1** and **Methods**). We generated worms that contain both *gfp* and *mcherry* transgenes, which are integrated into different genomic locations on different chromosomes but are constructed using similar promoters and 3’ UTRs sequences (See **Fig. 2A**). For these experiments, we selected worms in which the *mcherry* transgene is still expressed and varies in the population (and is not completely silenced) and exposed their progeny – the P0 generation – to anti-*gfp* RNAi. We then examined whether the level of spontaneous endogenous small RNA-mediated silencing of *mcherry* in the parental generation, which is exposed to the dsRNA trigger, could predict the fate of the dsRNA-initiated RNAi inheritance against the other gene, *gfp* (**Fig. 2B to C**). We found that P0 mothers which spontaneously silence the *mcherry* transgene trigger stronger and more penetrant heritable *gfp*-silencing responses across generations when exposed to anti-*gfp* dsRNA (**Fig. 2D**). Thus, the two silencing phenomena are coordinated, and their observed variability is shared. We conclude that each mother assumes a general, non-gene-target-specific “small RNA inheritance state”. Importantly, while the spontaneous silencing of the *mcherry* transgene predicted the strength of the dsRNA-triggered anti-*gfp* RNAi inheritance, the expression of the two transgenes in the absence of anti-*gfp* dsRNA-induced RNAi was not positively correlated (**Fig. S1**). Thus, internal inter-individual variability in initiating RNAi inheritance determines the fate of inherited responses that propagate and segregate across generations.

**Fig. 2.**
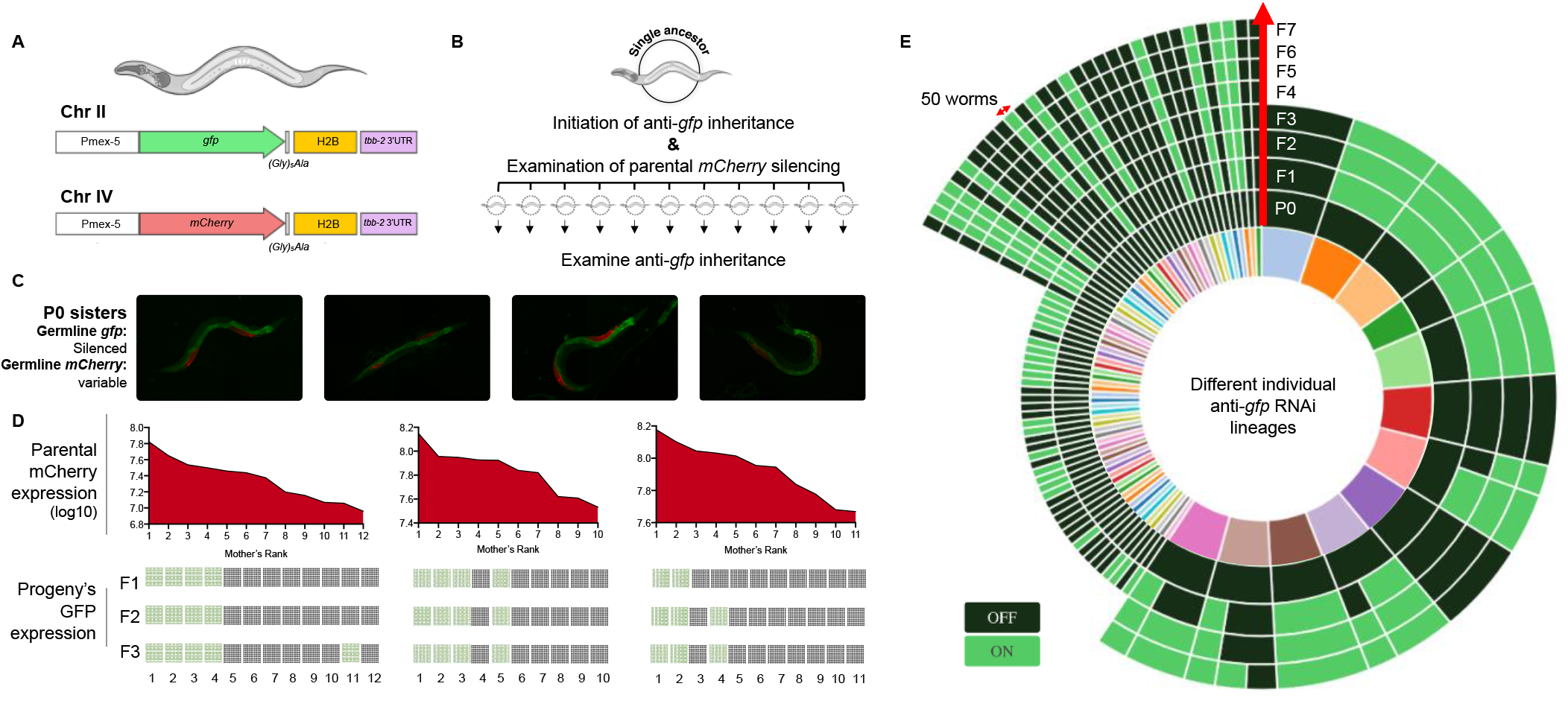
Inter-individual variability in *anti-gfp* small RNA inheritance can be predicted by the rate of spontaneous *mCherry* silencing at the parental generation. A. Transgene construction of genomically-integrated single-copies of *gfp* and *mCherry* transgenes. Both transgenes bear identical promoter and 3’ UTR sequences and were integrated using MosSCI into separate genomic loci and different chromosomes[42,43] (**Methods**). B. Experimental scheme: to examine the ability to predict inter-individual variability in silencing inheritance, worms which bear both transgenes were subjected to an anti-*gfp* RNAi treatment and were scored for spontaneous silencing of the *mcherry* transgene at the generation of exposure to RNAi. Experiments were done in triplicates, with each replicate of P0 mothers derived from one single ancestor. Inheritance of anti-*gfp* silencing response was scored at the next generations separately for each P0-derived lineage. C. Fluorescent microscope images of sister worms which are exposed to *anti-gfp* RNAi. While all mothers completely silence the *gfp* transgene upon exposure to RNAi, the *mcherry* transgene is spontaneously silenced to different degrees in sister worms. D. Matched scores of parental mCherry expression and inheritance of anti-*gfp* silencing across generations. Each individual P0 mother was examined for mCherry expression levels at the parental generation and ranked based on its relative expression compared to its sister worms. Presented are the log10 expression levels of mCherry for each mother, and the proportion of silencing (dark grey) vs. re-expressing (green) progeny in each lineage across generations. E. Visual summary of all anti-*gfp* RNAi inheritance lineages examined in this study (>20,000). Data are presented as inheritance “series”: each colored section of the inner circle represents one separate lineage, originating from an individual P0 mother. The outer circles represent each generation, and the scored silencing phenotype for the specified generation in each lineage. OFF: full silencing of *gfp.* ON: re-expression of GFP. The size of each section in the outermost circle corresponds to the number of worms examined in each lineage that were found to have the indicated silencing phenotype. Red arrow (↔) represents 50 worms.

Overall, and throughout the different experiments carried out in this study, we have examined inheritance in lineages composed of a total of >20,000 worms. To understand the transition rules of the silencing response across generations, we systematically analyzed these inheritance data and found a “hot hand” effect of transgenerational silencing: the more generations a heritable response lasted, the higher the chances that it will be transmitted to the next generation as well (46.5% of the worms kept silencing the *gfp* transgene at the transition from the 1st to the 2nd generation; 80.5% at the transition from the 2nd to the 3rd, 92.6%-100% starting from the 3rd generation. See summary visualization in **Fig. 2E**).

To characterize the differences in the RNA pools of worms that assume different RNAi inheritance states, we isolated four sister worms that derive from a single mother and allowed each of these worms to lay eggs for 6 hours on separate plates. We used a COPAS_TM_ “Worm Sorter” [28] to isolate worms that strongly express or spontaneously silence *mcherry* in each of these lineages (25 top and bottom percentile, see scheme in **Fig. 3A**). We sequenced small RNAs and mRNAs from the sorted worms to investigate what underlies the different internal inheritance states. The total levels of small RNA transcripts which derive from piRNA loci are *increased* in the sisters that silence the *mcherry* transgene (this transgene contains multiple piRNA-recognition sites [27], **Fig. 3B**). In contrast, we found that overall endogenous small RNA levels are reduced in worms which spontaneously silence the *mcherry* transgene (**Fig. 3B**). It is well-established that in worms different small RNA pathways compete over limited shared biosynthesis machineries [29]. It is therefore possible that the decrease in endogenous proteincoding-targeting small RNAs in *mcherry-*silencing worms makes more small RNA-processing machinery available for additional endogenous or exogenous siRNAs species. Together, these analyses suggest that sisters, despite being isogenic and deriving from the same mother, are equipped with a different arsenal of gene-silencing endogenous small RNAs.

**Fig. 3.**
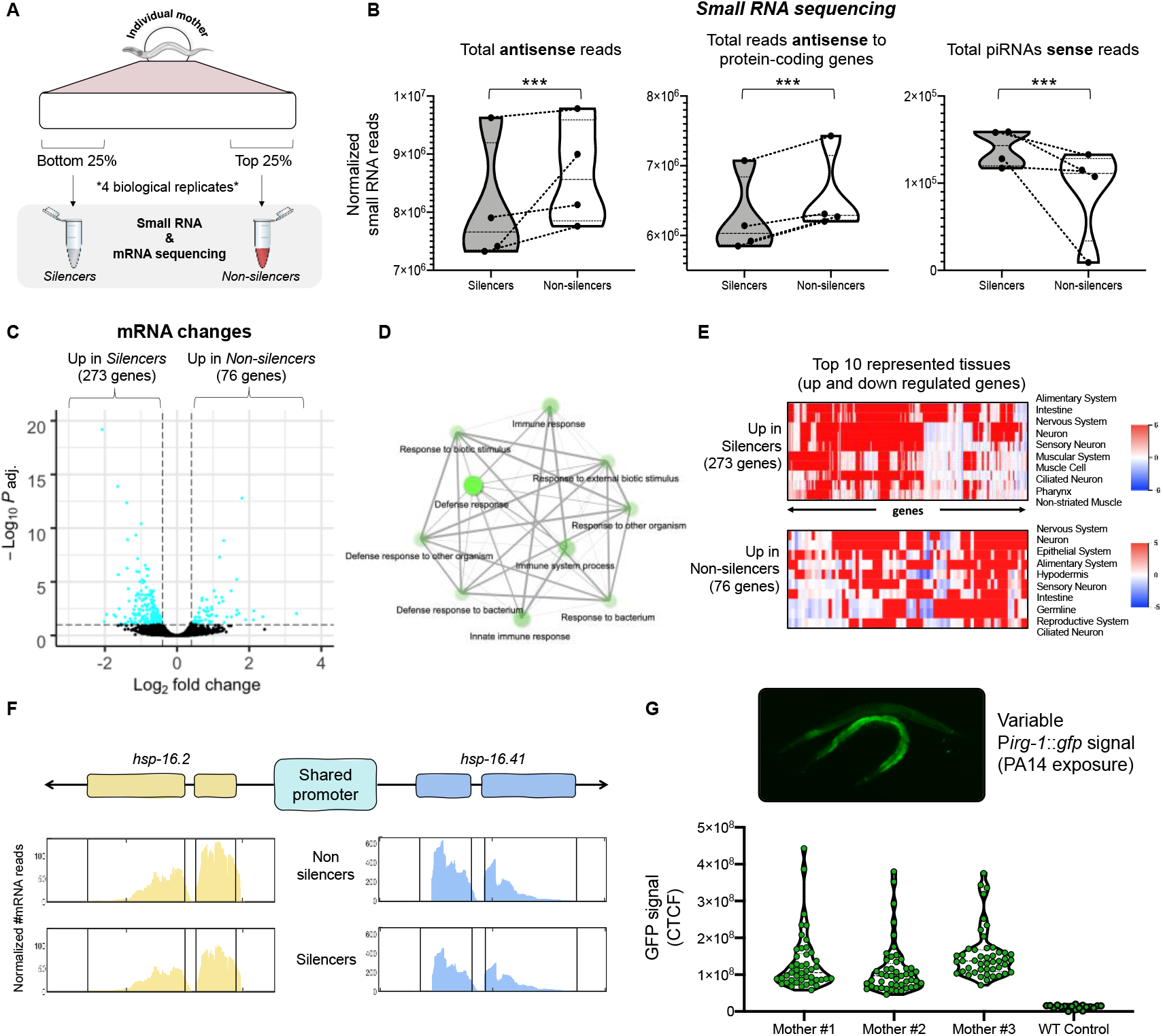
The transcriptomic changes that underlie the different inheritance states in isogenic sister worms. A. The experimental design for collecting tightly synchronized, isogenic sister worms bearing different inheritance states (**Methods**). B. Global changes in small RNA species in silencer vs. non-silencer worms (χ^2^ test). The Y-axis represents the normalized total number of reads for each indicated type of small RNA species. Each coupled measurement of progeny (silencer and non-silencer sisters) is marked with a dotted line. See **Methods** for full details regarding analyses. C. Volcano plot summarizing mRNA differences between silencer and non-silencer worms. D. Gene ontology (GO) enrichment[44] of genes which are differentially expressed between silencers and non-silencers, and the connection between the different enriched terms (top 10 enriched terms). Brighter nodes represent more significantly enriched gene sets. Bigger nodes represent larger gene sets for the specific GO term. Terms (nodes) are connected to each other if they share at least 20% of their genes. E. Tissue-enrichment analysis for genes which are up or down regulated between the two states of inheritance [31]. Red color represents higher predicted gene expression in the indicated tissue. F. The heat-shock protein genes *hsp-16.2* and *hsp-16.41* share the same promoter and their expression levels change together in the two different states of inheritance. Presented are normalized mRNA-seq reads along the two genes. Grey areas represent annotated exons locations. G. Inter-individual variability in the expression of an *irg-1* transcriptional reporter, one of the genes which were found to vary between the different inheritance states. Expression was measured in progeny of individual worms following 6 hours of exposure to the pathogenic bacterium *Pseudomonas aeruginosa* (PA14) (see **methods**).

We next examined the changes in mRNA expression levels in *mcherry* silencing and non-silencing sisters. Although we sequenced tightly synchronized isogenic progeny that were derived from a single mother, we found that 349 genes are differentially expressed between sisters which express or silence the *mcherry* transgene (**Fig. 3C**). Surprisingly, we found that the most significant enrichment in this set of genes is for immune and defense response processes (immune response, GO:0006955, FDR = 6.08E-09; defense response, GO:0006952, FDR = 2.08E-09, **Fig. 3D** and **Fig. S2**). This enrichment might echo the evolutionary origins of small RNAs as part of the worm’s immune system [30]. Interestingly, we found that these differentially expressed genes are enriched in neurons and in the alimentary system [31] (**Fig. 3E** and **Fig. S3**). The differentially expressed genes included multiple genes known to function together in specific regulatory pathways (see **Fig S2**, **Table S1**). Among the genes which vary between *mcherry*-silencers and non-silencers sisters, we identified *hsp-16.2*, a heat shock protein whose variable activation upon heat stress is a classic example for stochastic gene expression differences between isogenic worms. Non-genetic variable expression of HSP-16.2 was shown to predict the worm’s life span and stress resistance [2]. Interestingly, the promoter of *hsp-16.2* is shared (at the opposite orientation, see **Fig. 3F** and **Fig. S3**) with *hsp-16.41,* an additional Heat Shock Protein. We found that both *hsp-16.2* and *hsp-16.41* show similar transcriptional changes between the two states (**Fig. 3F** and **Fig. S3**), indicating that the observed variability arises from variation at the level of transcriptional regulation.

Among the different genes that show differential mRNA levels in our sequencing data we identified *irg-1* (Immune Response Gene-1). IRG-1 plays a critical role in immunity, and specifically in the response to pathogenic *pseudomonas* (PA14) [32]. We used a fluorescent reporter to quantify *irg-1* levels in individuals to validate the mRNA sequencing results. Although *irg-1* was never reported before to vary in isogenic populations, we found that *irg-1* indeed shows variable pathogen-induced induction levels in isogenic, synchronized, individuals (**Fig. 3G**).

As many of the genes that typify the two “heritable silencing states” have shared roles (e.g. immune regulation), and since we observed patterns of transcriptional regulation that vary between silencers and non-silencers (**Fig. 3F** and **Fig. S3**), we looked for regulators (e.g. transcription factors) that might synchronize the RNAi inheritance states. The existence of multiple Heat Shock Proteins which vary between the two states, and the unexpected enrichment for cuticle-structure genes pointed toward Heat Shock Factor-1 (HSF-1)-dependent regulation [33]. HSF-1 is a highly conserved transcription factor, a master regulator of proteostasis, whose activity is important under both stressful and non-stressful conditions [34]. HSF-1 was recently shown to be involved in transcriptome remodeling of non-coding RNAs’ pools in response to heat shock in *C. elegans* nematodes [35] and to account for cell-to-cell variation and phenotypic plasticity in budding yeast [36].

We found that the genes which are differentially expressed between the two inheritance states show a strong enrichment for HSF-1-regulated genes [33] (166/349 genes, 1.63 fold enrichment, p-value < 2.45e-13, hypergeometric test **Fig. 4A**), and that gene expression changes in the non-silencers worms resemble gene expression changes following knockdown of *hsf-1* [33] (**Fig. 4A**).

**Fig. 4.**
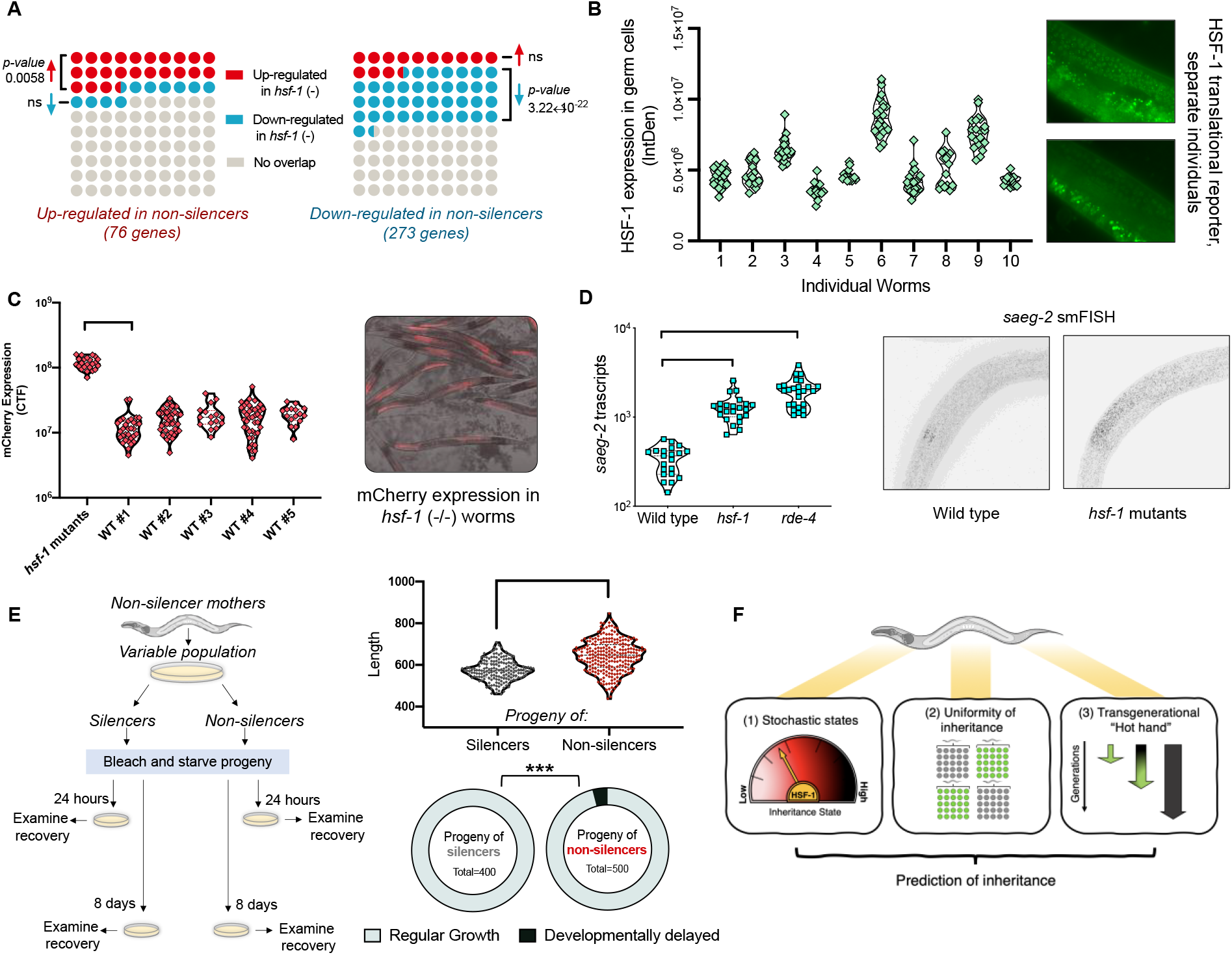
HSF-1 varies between different individuals and affects multiple inheritance processes. The inheritance states can predict the next generations’ response to starvation stress. A. Genes which are up and down regulated in the two different inheritance states show enrichment for HSF-1-regulated genes. Specifically, transcriptional changes in non-silencers resemble transcriptional changes in *hsf-1* mutant worms (hypergeometric test). B. Translational reporter of the worm’s HSF-1 protein reveals inter-individual variation in the formation of HSF-1 stress granules in the germline. Each dot represents GFP levels in one syncytial nucleus. Data were collected from ten separate and tightly synchronized individual worms (X axis). C. Variability in spontaneous silencing of the *mcherry* transgene is abolished in worms which possess a reduction-of-function mutation in *hsf-1 (sy441).* Each dot represents a measurement of total fluorescence in one individual worm (F test). D. Quantifications of *saeg-2* transcripts using smFISH in wild type, *hsf-1 (sy441)* and *rde-4 (ne299)* worms. Each dot represents one quantified worm. P-values were determined by Kruskal-Wallis test with Dunn’s post hoc correction for multiple comparison and asterisks represent p-values in comparison to wild type. **** represents p-value < 10^-4^. E. The worms’ response to starvation stress can be predicted by examination of the inheritance state at the parental generation, prior to starvation (see **Methods**). Progeny of non-silencer worms show pronounced variability and developmental delay following recovery from 8 days of starvation. Top panel: length measurements [45] 60 hours after transferring the worms to plates with food (F test). Bottom panel: frequency of developmentally delayed worms (arrest as L1/L2) 48 hours after transferring the worms to plates with food (χ^2^ test). F. Rules for the segregation of heritable small RNA responses across generations: (1) each mother assumes a stochastic “inheritance state”, which dictates the potency of the small RNA responses that this worm would transmit to its descendants, (2) within every generation heritable silencing responses are distributed uniformly among the progeny of the same RNAi-exposed mother, (3) The more generations silencing responses last, the chance that silencing would persist increases, according to the “hot hand” principle.

To directly test the involvement of HSF-1 in generating variable inheritance states, we first examined whether individual worms display variability in HSF-1’s function. We did not detect significant inter-individual variability in *hsf-1* mRNA levels, in accordance with previous observations of HSF-1’s activity being regulated at the post-translational level [37]. Specifically, it was recently revealed that in *C. elegans* HSF-1’s activity corresponds to the formation of stress granules inside the nucleus [38]. By using a single-copy integrated fluorescent GFP translational reporter, we found that the number of HSF-1 granules and HSF-1’s expression intensity vary in the germline syncytium nuclei of isogenic and tightly synchronized sister worms, even in the absence of any external stress or heat-shock stimuli (**Fig. 4B**).

To investigate if HSF-1 plays a role in constituting different inheritance states, we tested how changes in HSF-1’s activity affect small RNA-mediated heritable silencing. We examined worms which bear a reduction-of-function mutation in *hsf-1 (sy441)* and measured the state of their RNAi inheritance machinery. We quantified how the rate and variability of spontaneous *mcherry* silencing changes in *hsf-1* reduction-of-function mutants. Strikingly, we found that in these mutants the differences in heritable silencing between isogenic sisters were canceled, namely, all the worms exhibited a uniform expression of the *mcherry* transgene (**Fig. 4C**). We then examined whether HSF-1’s activity can also affect *endogenous* transgenerational silencing functions in the worms. We recently reported that the *saeg-2* gene is strongly silenced in the germline by RDE-4-dependent endo-siRNAs, and that neuronally-transcribed small RNAs control this silencing transgenerationally [22]. Using single-molecule fluorescence in situ hybridization (smFISH), we found that worms which bear a reduction-of-function mutation in *hsf-1* de-silence *saeg-2,* similarly to the de-silencing that is observed in mutants devoid of *saeg-2*-targeting siRNAs [22] (**Fig. 4D**). These results, together with previous evidence that HSF-1 sculpts the pools of different endogenous small RNAs [35], suggest that HSF-1 affects multiple, separate transgenerational small RNA processes, and as such, can coordinate the broad small RNA inheritance “states”.

Survival during long periods of starvation requires HSF-1 [39]. In the absence of food, worms arrest their development at the first larval stage; Starvation alters the transgenerational pool of heritable small RNAs [6,24], and leads to multiple phenotypic changes in the progeny, such as increased lifespan [6], reduced fecundity and increased heat resistance [40]. Interestingly, isogenic worms produce different transgenerational responses following starvation [40]. We hypothesized that by monitoring the inheritance states of the parents, we can predict the transgenerational phenotypic responses to starvation. We found that starved progeny of mothers that silence the *mcherry* transgene did not show pronounced variability in recovery time following starvation as measured by size distribution and the presence of L1/2-arrested worms in the population (**Fig. 4E**). In striking contrast, progeny of mothers that did not silence the *mcherry* transgene exhibited high variability in their recovery time following starvation and higher rates of developmental delay (**Fig. 4E**). Overall, these results indicate that the mother’s “inheritance state” predicts multiple heritable small RNA-related phenotypes: transgenerational dsRNA-induced silencing, spontaneous transgene-silencing, and the inherited outcomes of environmental stress-relevant responses.

In summary, we find that the propagation of small RNA-encoded epigenetic information across generations can be explained using simple rules (Fig. 4F). Variation is the raw material of evolution, and while small RNAs could increase inter-individual variability, and some somatic processes in the worm can control DNA changes in the worm’s germline [41], it is still unclear if and how epigenetic inheritance contributes to the process of evolution. Our results suggest that similarly to the mechanisms which evolved to generate genetic variation (e.g. random assortment of chromosomes and recombination), an innate mechanism gives rise to variability in small RNA inheritance.

## Acknowledgments

We thank all the Oded Rechavi lab members for comments and discussions. Special thanks to Sarit Anava, Ekaterina Star, Hila Gingold and Guy Teichman for experimental and visualization assistance. Some strains were provided by the CGC, which is funded by NIH Office of Research Infrastructure Programs (P40 OD010440). We thank Eric Miska and Julie Ahringer for kindly providing us with strains. L.H.Z. is thankful to the Clore Foundation. O.R. is thankful to the Adelis Foundation grant #0604916191. This work was funded by ERC grants #335624 and # 819151. As this is a short article, we apologize that we could not reference many other important studies on the mechanisms of RNAi and small RNA inheritance, especially experiments that were conducted with organisms other than *C.elegans.*

## Methods

### Nematodes’ growth and maintenance

Standard culture techniques were used to maintain the worms. Worms were grown on Nematode

Growth Medium (NGM) plates and fed with *Escherichia coli* OP50 bacteria. Strains maintenance and experiments were performed at 20°C. The worms were kept fed for at least five generations before the beginning of each experiment. Extreme care was taken to avoid contamination or starvation. Contaminated plates were discarded from the analysis.

### Strains

The following *C. elegans* strains were used in this study: N2, SX1263 (mjIs134 [Pmex-5::gfp::(Gly)5Ala/his-58/tbb-2 3’UTR; cb-unc-119(+)] II), JA1527 (weSi14 [Pmex-5::mCherry::(Gly)5Ala/his-58/tbb-2 3’UTR; cb-unc-119(+)] IV)[42], PS3551 *(hsf-1(sy441)),* OG532 (drSi13 [Phsf-1::hsf-1::gfp::unc-54 3’UTR + Cbr-unc-119(+)] II) - outcrossed to generate wild-type background, AU133 (agIs17 [Pmyo-2::mCherry + Pirg-1::GFP] IV), BFF12 *(rde-4 (ne299)).*

### dsRNA administration

The standard assay for RNAi by feeding was carried out as previously described[46]: HT115 bacteria that transcribe dsRNA targeting *gfp* were grown in Carbenicillin-containing LB and were than seeded on NGM plates that contain Carbenicillin (100 μg/ml) and IPTG (1 mM). The plates were seeded with bacteria 24 hours prior to their use.

### Lineages experiments

*Diversifying anti-gfp RNAi lineages* (**Fig. 1A**):

An individual worm (day 2 of adulthood) was placed on a plate containing anti-*gfp* producing bacteria and was given 2-3 hours to lay eggs (the P0 generation). We then let the eggs hatch, and the worms grow until they become young adults. Four P0 worms were then transferred to plates containing regular bacteria (OP50) and were allowed to lay eggs (the F1 generation) for 6-8 hours. The P0 mothers were then removed and pictured for their silencing phenotype (all P0 mothers exhibited complete silencing).

The eggs (F1 generation) were then allowed to hatch, and the worms grew until reaching day 2 of adulthood. On day 2, we randomly selected 3 F1 worms from each separate P0-derived group and transferred each of them to a new plate for 12-16 hours of egg laying. We then pictured the entire F1 population and the three randomly selected F1 mothers. We repeated this procedure in the transmission between the F2 and F3 generation. In the F3 generation all 36 progeny groups were pictured. Across all generations, each group of progenies was constructed of 25-90 worms, with an average of 43 worms per group (median = 41.5).

(4 P0s x 3 F1s each x 3F2s each = 36 F3 groups)

*Multiple separate anti-gfp RNAi P0 mothers* (**Fig. 1B**):

To control for differences in *in utero* exposure of both the P0 and the F1 generation, we exposed the ancestor worm (which will lay the P0 generation) to the RNAi signal from the beginning of the L4 stage (before the creation of eggs), and allowed the P0 worms to “recover” on control plated before laying the F1 generation that will be examined in the experiments. Namely: an individual L4 worm was placed on a plate containing anti-*gfp* producing bacteria until it reached day 2 of adulthood. It was then transferred to a new anti-*gfp* plate and was allowed to lay eggs for 6-8 hours (the P0 generation). The eggs were allowed to hatch, and the worms grew until they reached day 1-2 of adulthood. The P0 mothers were transferred to regular plates for 4 hours (during which the fertilized eggs that were directly exposed to the RNAi signal should have all been laid). The P0 mothers were then transferred to separate plate (1 worm per plate) to lay eggs (the F1 generation) for 6-8 hours, and then removed and pictured for the silencing phenotype. The eggs (F1 generation) hatched and the worm grew until day 1 of adulthood, in which they were scored for their silencing phenotype.

*Diversifying mcherry stochastic silencing lineages experiments* (**Fig. S1**). Five mCherry-expressing worms were chosen to create the five different lineages. Similar to anti-*gfp* lineages experiments, three worms were randomly selected from the progeny to generate the next generation. This procedure repeated in each examined generation (until F4).

*Coupled mcherry (stochastic silencing) and *gfp* (dsRNA-induced) lineages* (**Fig. 2A to D**) For each of the three replicates, a single expressing adult (day 2) worm was chosen and was allowed to lay eggs (the P0 generation) on an *anti-gfp* RNAi plate for 6-8 hours. The eggs were allowed to hatch, and the worms grew until they reached adulthood. The P0 worms were then transferred to regular plates for 4 hours of “recovery” egg laying (to minimize differences in *in utero* exposure of the eggs to the dsRNA signal). The P0 mothers were then transferred to separate plates – one P0 worm per plate – and allowed to lay eggs (the F1 generation) for 6-8 hours. The mothers were then removed and pictured for the anti-*gfp* silencing phenotype and for their *mcherry* silencing phenotype. The eggs were allowed to hatch, and worms grew until they reached day 2 of adulthood, at which each group was pictured for the anti-*gfp* silencing phenotype and one worm was transferred to a new plate to lay the next generation (12-16 hours of egg laying). This procedure repeated itself in each examined generation.

### Handling of the JA1527 strain

*(bearing a stochastically silenced mcherry transgene)*

The JA1527 strain was kindly received from the Julie Ahringer lab (University of Cambridge). After 6 outcrosses rounds, rapid silencing of the *mcherry* transgene was initiated in the population. To avoid a complete “drift” of *mcherry* silencing in the population we either (1) kept large stocks of frozen expressing JA1527 worms and thawed the worms a couple of weeks prior to the initiation of experiments or (2) transferred the worms to 25 °C for a couple of generations, transferred them back to 20°C and selected multiple expressing worms to create the next generations. We let these worms revive for at least 3 generations before the initiation of experiments.

It is highly recommended to avoid repetitive selection of expressing worms (tens of generations of selection), as it seems to drastically change RNAi sensitivity in the population.

### RNA and small RNA sequencing experiments

*Collecting worms for sequencing:* four mCherry-expressing worms were allowed to lay eggs in separate plates (one worm per plate) for 8 hours. The eggs were allowed to hatch and grew until they reached adulthood. Each group of progenies (isogenic sisters) was then washed with M9 buffer into a 1.5ml Eppendorf tube, followed by 3-4 M9 washes, in order to remove any residual bacteria. Each group was then examined in COPAS_TM_ Biosort (Union Biometrica) and sorted to get the top and bottom 25% of mCherry expressing worms in each group. TRIzol^®^ (Life Technologies) was then added to each sorted tube, and the tubes were immediately transferred to −80°C until RNA extraction procedure.

*RNA extraction:* RNA extraction was performed as previously described[22].

*Small RNA libraries*: total RNA samples were treated with tobacco acid pyrophosphatase (TAP, Epicenter), to ensure 5’ monophosphate-independent capturing of small RNAs. Libraries were prepared using the NEBNext^®^ Small RNA Library Prep Set for Illumina^®^ according to the manufacturer’s protocol. The resulting cDNAs were separated on a 4% agarose E-Gel (Invitrogen, Life Technologies), and the 140–160 nt length species were selected. cDNA was purified using the MinElute Gel Extraction kit (QIAGEN). Libraries were sequenced using an Illumina HiSeq 2500 instrument.

*mRNA libraries:* cDNA was generated using SMART-Seq v4 Ultra Low Input RNA (Takara) and libraries were prepared using Nextera XT DNA Sample Preparation Kit (Illumina). Libraries were sequenced using an Illumina MiniSeq instrument.

### Sequencing analyses

*Small RNA libraries.* Small RNA libraries were processed and analyzed as recently described[24]. Normalization of the total number of reads in each sample, and the total number of reads which align to the different types of genomic features was generated based on the SizeFactor normalization provided by the DESeq2 package (the median ratio method). To omit effects of PCR amplifications and additional sources of variability which are driven by sequencing artifacts, Chi square test was performed on DESeq2-size-factor normalized reads and the statistic value of the test was normalized to the same size factors.

*mRNA libraries*. mRNA libraries were first assessed for quality using the FastQC tool[47] and were then aligned to ce11 version of the genome using bowtie 2[48], using the command: *bowtie2--sensitive-local -x ce11Reference-UmRNA_sample.fastq.gz > Alignment.sam*

The aligned reads were then counted using the python-based script HTSeq-count[49] and the Ensembl-provided gff file (release-95), using the following command:

*HTSeq.scripts.count --stranded=yes --mode=union Alignment.sam ce11WBcel235.94.gtf > Counts.txt*

The samples were then compared for differential expression using the R package DESeq2[50], and a “patient”-based comparison, to directly compare between each pairs of silencer and non-silencer sisters.

### PA14 exposure experiments

Corresponds to **Fig. 3G**: worms were exposed to PA14 as previously described[32]. Adult (day 2) AU133 and wild type worms were allowed to lay eggs in separate OP50 plates (single worm per plate) for 10 hours. The eggs hatched and the worms grew until they reached adulthood. The worms were then placed on PA14-containing plates for 8 hours. After 8 hours the worms were removed and pictured for *Pirg-1*::GFP expression.

### HSF-1 translational reporter experiments

Corresponds to **Fig. 4B**: Worms were first outcrossed with wild-type (N2) worms to generate worms which bear genomically integrated Phsf-1::hsf-1::gfp::unc-54 3’UTR construct in a wildtype background. A single worm was allowed to lay eggs for 12 hours. L4 worms were selected from the progeny to further facilitate synchronization between the different individuals and were examined for the HSF-1 translational reporter expression 24 hours later.

### smFISH experiments

smFISH experiments and analyses were performed as previously described[22] using the same probe set.

### Starvation experiments

Corresponds to **Fig. 4E**: Multiple mCherry-expressing adult worms were allowed to lay eggs over-night. The eggs hatched and the worms grew, until they reached day 2 of adulthood. The population was separated into silencers and non-silencers groups based on mCherry fluorescence. Each group was bleached to achieve clean batches of eggs. The eggs were transferred to unseeded Nematode Growth Medium (NGM) plates for either 24 hours of 8 days. After the indicated starvation period, worms were washed and transferred to plates with food (OP50 bacteria) and were scored for size and developmental delay.

*Scoring of developmentally delayed worms:* After 48 hours of recovery, the worms were examined for the existence of developmentally delayed worms (arrested at the L1/L2 stage). The number of all worms and the developmentally delayed worms in each condition was counted. The investigators were blinded to identity of the groups during their examination.

*Measuring worms’ size using WorMachine:* After 60 hours of recovery, each group of worms was washed, paralyzed, pictured and then analyzed as previously described[45], using the WorMachine software.

### Microscopy

We used an Olympus BX63 microscope for fluorescence microscopy assays. Unless otherwise noted experiments were pictured using a 10X objective lens, and an exposure time of 750ms. Generally, worms were picked and transferred to 2% agarose padded microscope slides containing drops of tetramisole to generate paralysis, covered with a glass cover slip, and pictured after 2-5 minutes.

*Measuring anti-gfp RNAi inheritance.* GFP silencing and inheritance was scored using a binary system: no expression (OFF) or any level of expression (ON).

Measuring *mCherry’s* stochastic silencing and *irg-1* reporter. Using *Fiji,* we measured the integrated density of the whole worm, based on its contour, as well as multiple background measurements per picture. The corrected total fluorescence (CTF) of each worm was calculated as {Integrated Density – (Area of selected worm X Mean fluorescence of background readings)}. *Measuring hsf-1 reporter.* The syncytial germline of the worms was pictured using 40X objective lens. The GFP fluorescence in nuclei of each worm were then measured using *Fiji*.

### Statistics and visualization

All statistical analyses were performed in R (version 3.6.1) or Graphpad Prism (version 8.3.0). Statistical tests were performed as two-sided tests, unless inapplicable. Chi-square test and hypergeometric tests were performed using online tools. Graphs were created in R and Graphpad Prism.

**Extended Data Figure 1:**
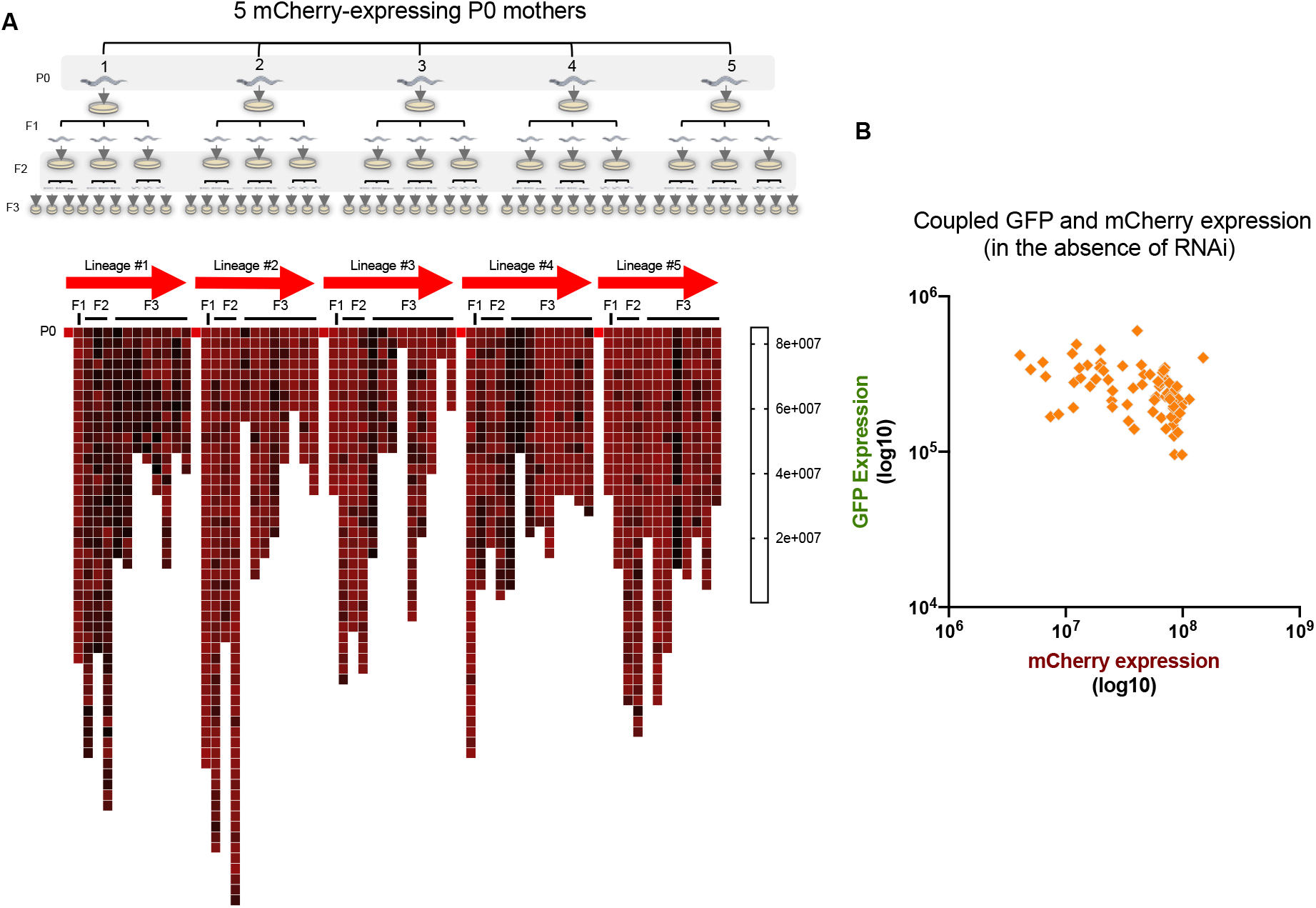
Stochastic silencing of the *mcherry* transgene across lineages, and in comparison to GFP expression in the absence of anti-*gfp* RNAi. **A.** Top panel: scheme of examination of *mcherry* stochastic silencing across lineages (see **Methods**). Bottom: mCherry expression levels in each examined worm across the different lineages. Scale represent high (red) and low (dark) mCherry expression levels. Each square represents a measurement of a single worm. Few groups were omitted from the analysis due to contamination or premature death. B. Paired measurements of GFP and mCherry expression in the absence of anti-*gfp* RNAi treatment in worms which bear the two transgenes. Each dot represents the GFP and the mCherry measurements of a single worm. Slight negative correlation is observed between the two expression levels (simple linear regression. Slope = −0.001155, p-value < 0.0005)

**Extended Data Figure 2:**
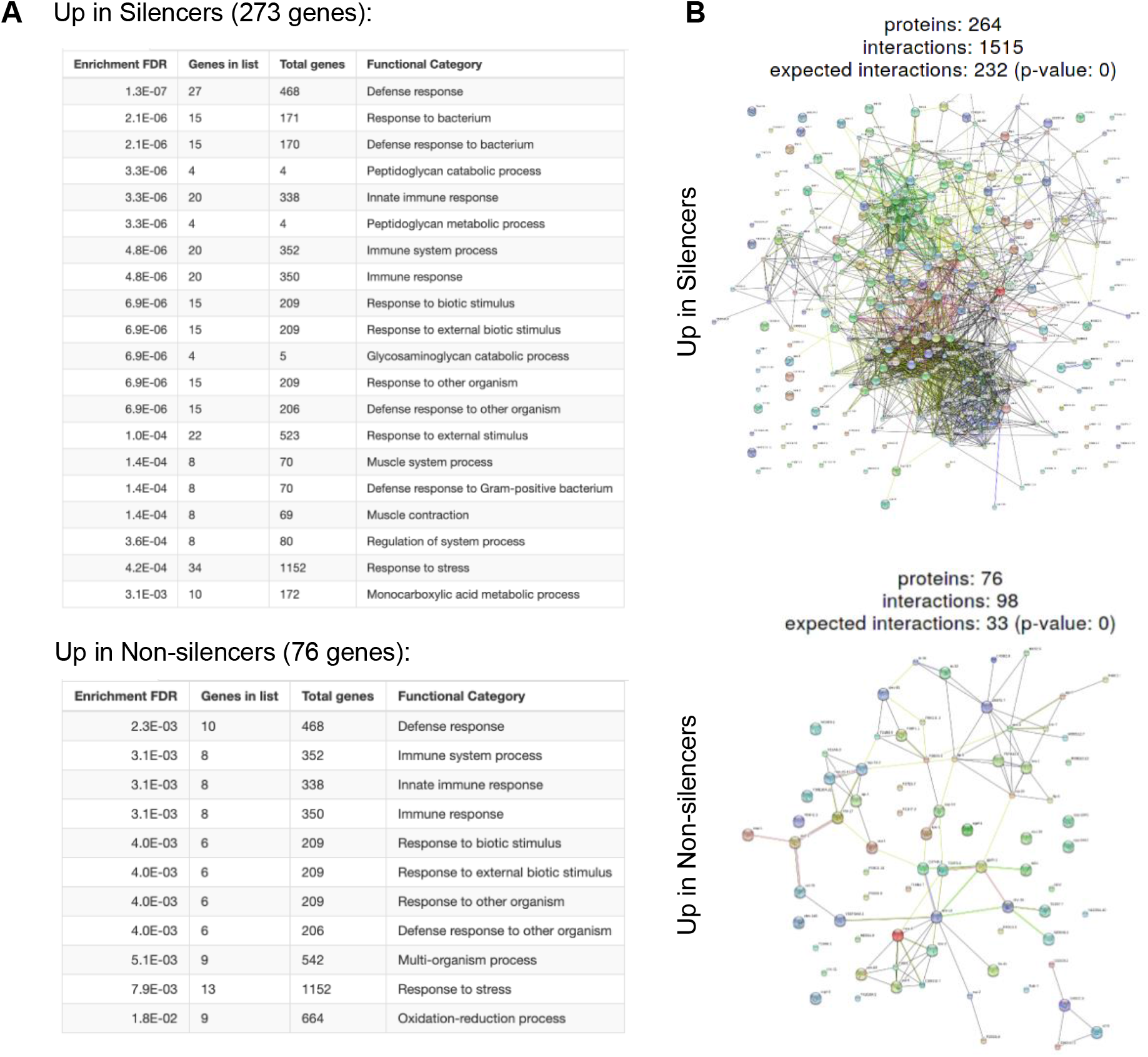
GO term enrichment and STRING representation of genes which are differentially expressed between the two inheritance states. **A.** Gene ontology (GO) for genes which are up regulated in silencer and non-silencer worms. Presented are the top 20 terms for each dataset, which have an FDR enrichment value of less than 0.05 (only 11 terms pass this criterion in the dataset of genes which are up regulated in Non-silencers). B. STRING representation of the protein-protein interactions between the genes which are differentially expressed in the different states of inheritance. Indicated is the expected number of interactions, the observed number, and a significance value for enrichment of interactions (Fisher’s exact test followed by a correction for multiple testing). Both panels were created using ShinyGO v0.61[44].

**Extended Data Figure 3:**
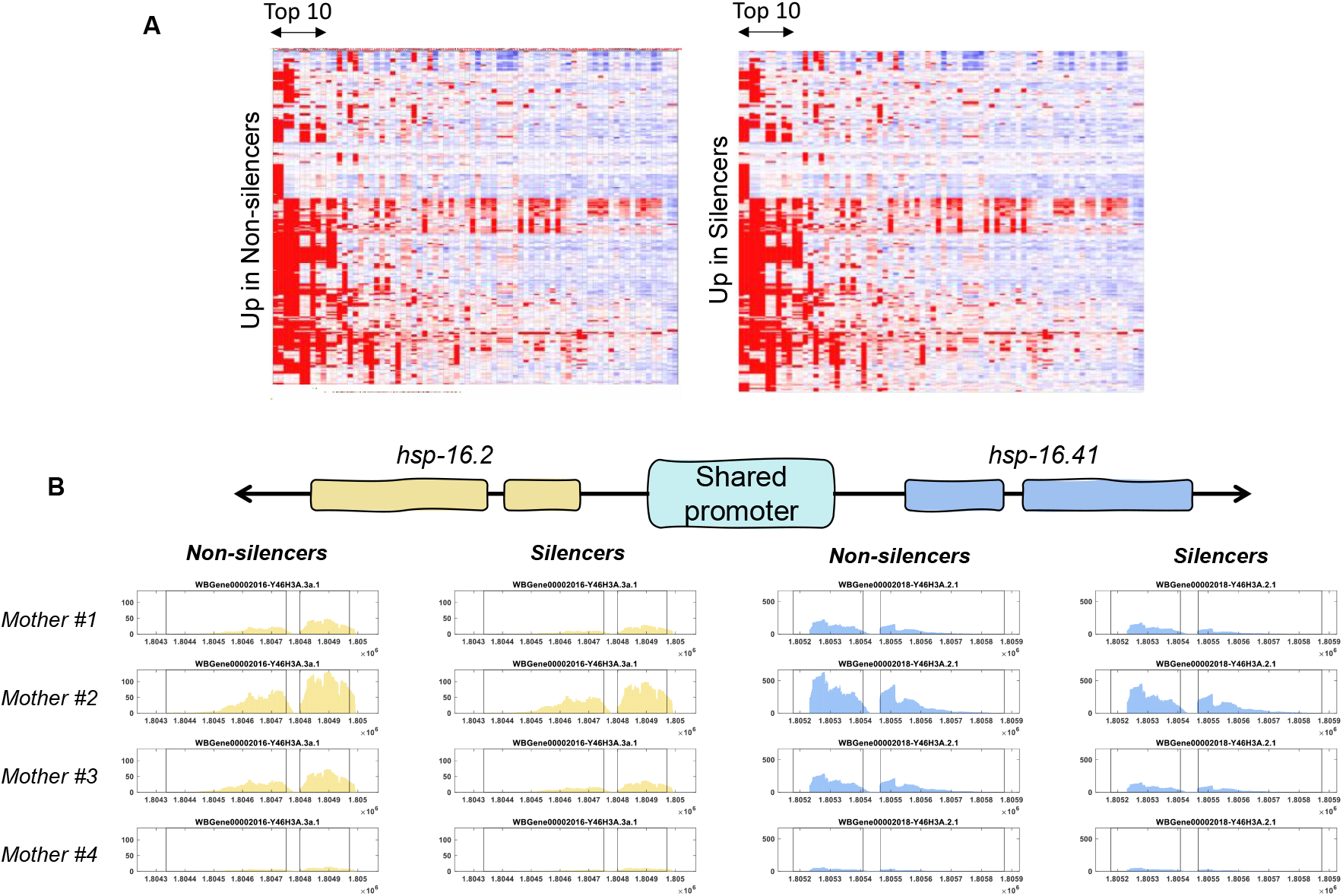
Tissue expression prediction of genes which change their expression in the different inheritance states, and co-regulation of the *hsp-16.2* and *hsp-16.41* genes, which share a promoter. **A.** Full visualization of tissue expression prediction of the genes which are up regulated in Non-silencers (left) and Silencers (right). Columns represent different tissues, rows represent the different genes in each dataset[31]. B. Normalized mRNA-seq reads aligning to *hsp-16.2* and *hsp-16.41* which share their promoters. Presented are the normalized reads for each of the four groups of progenies which were sorted into Silencer and Non-silencer worms.

## References and Notes

1. Stern S, Kirst C, Bargmann CI (2017) Neuromodulatory Control of Long-Term Behavioral Patterns and Individuality across Development. Cell 171: 1649–1662.e10.

2. Rea SL, Wu D, Cypser JR, Vaupel JW, Johnson TE (2005) A stress-sensitive reporter predicts longevity in isogenic populations of Caenorhabditis elegans. Nat Genet 37: 894–898.

3. Burga A, Casanueva MO, Lehner B (2011) Predicting mutation outcome from early stochastic variation in genetic interaction partners. Nature 480: 250–253.

4. Casanueva MO, Burga A, Lehner B (2012) Fitness Trade-Offs and Environmentally Induced Mutation Buffering in Isogenic C. elegans. Science (80-) 335: 82–85.

5. Perez MF, Francesconi M, Hidalgo-Carcedo C, Lehner B (2017) Maternal age generates phenotypic variation in Caenorhabditis elegans. Nature 552: 106–109.

6. Rechavi O, Houri-Ze’evi L, Anava S, Goh WSS, Kerk SY, Hannon GJ, Hobert O (2014) Starvation-induced transgenerational inheritance of small RNAs in C. elegans. Cell 158: 277–287.

7. Bazopoulou D, Knoefler D, Zheng Y, Ulrich K, Oleson BJ, Xie L, Kim M, Kaufmann A, Lee Y-T, Dou Y, et al. (2019) Developmental ROS individualizes organismal stress resistance and lifespan. Nature 1–5.

8. Houri-Ze’evi L, Korem Y, Sheftel H, Faigenbloom L, Toker IA, Dagan Y, Awad L, Degani L, Alon U, Rechavi O (2016) A Tunable Mechanism Determines the Duration of the Transgenerational Small RNA Inheritance in C. elegans. Cell 165:.

9. Alcazar RM, Lin R, Fire AZ (2008) Transmission dynamics of heritable silencing induced by double-stranded RNA in Caenorhabditis elegans. Genetics 180: 1275–1288.

10. Vastenhouw NL, Brunschwig K, Okihara KL, Müller F, Tijsterman M, Plasterk RHA (2006) Gene expression: long-term gene silencing by RNAi. Nature 442: 882.

11. Shirayama M, Seth M, Lee H-C, Gu W, Ishidate T, Conte D, Mello CC (2012) piRNAs initiate an epigenetic memory of nonself RNA in the C. elegans germline. Cell 150: 65–77.

12. Gu SG, Pak J, Guang S, Maniar JM, Kennedy S, Fire A (2012) Amplification of siRNA in Caenorhabditis elegans generates a transgenerational sequence-targeted histone H3 lysine 9 methylation footprint. Nat Genet 44: 157–164.

13. Luteijn MJ, van Bergeijk P, Kaaij LJT, Almeida MV, Roovers EF, Berezikov E, Ketting RF (2012) Extremely stable Piwi-induced gene silencing in Caenorhabditis elegans. EMBO J 31: 3422–3430.

14. Perales R, Pagano D, Wan G, Fields BD, Saltzman AL, Kennedy SG (2018) Transgenerational Epigenetic Inheritance Is Negatively Regulated by the HERI-1 Chromodomain Protein. Genetics 210: 1287–1299.

15. Ashe A, Sapetschnig A, Weick E-M, Mitchell J, Bagijn MP, Cording AC, Doebley A-L, Goldstein LD, Lehrbach NJ, Le Pen J, et al. (2012) piRNAs can trigger a multigenerational epigenetic memory in the germline of C. elegans. Cell 150: 88–99.

16. Moore RS, Kaletsky R, Murphy CT (2019) Piwi/PRG-1 Argonaute and TGF-β Mediate Transgenerational Learned Pathogenic Avoidance. Cell 177: 1827–1841.e12.

17. Rechavi O, Minevich G, Hobert O (2011) Transgenerational Inheritance of an Acquired Small RNA-Based Antiviral Response in C. elegans. Cell 147: 1248–1256.

18. Schott D, Yanai I, Hunter CP (2014) Natural RNA interference directs a heritable response to the environment. Sci Rep 4: 7387.

19. Ni JZ, Kalinava N, Chen E, Huang A, Trinh T, Gu SG (2016) A transgenerational role of the germline nuclear RNAi pathway in repressing heat stress-induced transcriptional activation in C. elegans. Epigenetics Chromatin 9: 3.

20. Hall SE, Chirn G-W, Lau NC, Sengupta P (2013) RNAi pathways contribute to developmental history-dependent phenotypic plasticity in C. elegans. RNA 19: 306–319.

21. Sims JR, Ow MC, Nishiguchi MA, Kim K, Sengupta P, Hall SE (2016) Developmental programming modulates olfactory behavior in C. elegans via endogenous RNAi pathways. Elife 5:.

22. Posner R, Toker IA, Antonova O, Star E, Anava S, Azmon E, Hendricks M, Bracha S, Gingold H, Rechavi O (2019) Neuronal Small RNAs Control Behavior Transgenerationally. Cell 177: 1814–1826.e15.

23. Vasale JJ, Gu W, Thivierge C, Batista PJ, Claycomb JM, Youngman EM, Duchaine TF, Mello CC, Conte D (2010) Sequential rounds of RNA-dependent RNA transcription drive endogenous small-RNA biogenesis in the ERGO-1/Argonaute pathway. Proc Natl Acad Sci USA 107: 3582–3587.

24. Houri-Ze’evi L, Teichman G, Gingold H, Rechavi O (2019) Stress Resets Transgenerational Small RNA Inheritance. bioRxiv 669051.

25. Lev I, Seroussi U, Gingold H, Bril R, Anava S, Rechavi O (2017) MET-2-Dependent H3K9 Methylation Suppresses Transgenerational Small RNA Inheritance. Curr Biol 27: 1138–1147.

26. Lev I, Toker IA, Mor Y, Nitzan A, Weintraub G, Antonova O, Bhonkar O, Ben Shushan I, Seroussi U, Claycomb JM, et al. (2019) Germ Granules Govern Small RNA Inheritance. Curr Biol 29: 2880–2891.e4.

27. Zhang D, Tu S, Stubna M, Wu W-S, Huang W-C, Weng Z, Lee H-C (2018) The piRNA targeting rules and the resistance to piRNA silencing in endogenous genes. Science 359: 587–592.

28. Pulak R (2006) Techniques for Analysis, Sorting, and Dispensing of *C. elegans* on the COPAS™ Flow-Sorting System. In, C. elegans pp 275–286. Humana Press, New Jersey.

29. Zhuang JJ, Hunter CP (2012) The Influence of Competition Among C. elegans Small RNA Pathways on Development. Genes (Basel) 3:.

30. Yan N, Chen ZJ (2012) Intrinsic antiviral immunity. Nat Immunol 13: 214–222.

31. Kaletsky R, Yao V, Williams A, Runnels AM, Tadych A, Zhou S, Troyanskaya OG, Murphy CT (2018) Transcriptome analysis of adult Caenorhabditis elegans cells reveals tissue-specific gene and isoform expression. PLOS Genet 14: e1007559.

32. Dunbar TL, Yan Z, Balla KM, Smelkinson MG, Troemel ER (2012) C. elegans detects pathogen-induced translational inhibition to activate immune signaling. Cell Host Microbe 11: 375–386.

33. Brunquell J, Morris S, Lu Y, Cheng F, Westerheide SD (2016) The genome-wide role of HSF-1 in the regulation of gene expression in Caenorhabditis elegans. BMC Genomics 17: 559.

34. Li J, Chauve L, Phelps G, Brielmann RM, Morimoto RI (2016) E2F coregulates an essential HSF developmental program that is distinct from the heat-shock response. Genes Dev 30: 2062–2075.

35. Schreiner WP, Pagliuso DC, Garrigues JM, Chen JS, Aalto AP, Pasquinelli AE (2019) Remodeling of the Caenorhabditis elegans non-coding RNA transcriptome by heat shock. Nucleic Acids Res 47: 9829–9841.

36. Zheng X, Beyzavi A, Krakowiak J, Patel N, Khalil AS, Pincus D (2018) Hsf1 Phosphorylation Generates Cell-to-Cell Variation in Hsp90 Levels and Promotes Phenotypic Plasticity. Cell Rep 22: 3099–3106.

37. Huang C, Wu J, Xu L, Wang J, Chen Z, Yang R (2018) Regulation of HSF1 protein stabilization: An updated review. Eur J Pharmacol 822: 69–77.

38. Morton EA, Lamitina T (2013) Caenorhabditis elegans HSF-1 is an essential nuclear protein that forms stress granule-like structures following heat shock. Aging Cell 12: 112–120.

39. Roux AE, Langhans K, Huynh W, Kenyon C (2016) Reversible Age-Related Phenotypes Induced during Larval Quiescence in C. elegans. Cell Metab 23: 1113–1126.

40. Jobson MA, Jordan JM, Sandrof MA, Hibshman JD, Lennox AL, Baugh LR (2015) Transgenerational Effects of Early Life Starvation on Growth, Reproduction and Stress Resistance in Caenorhabditis elegans. Genetics 201: 201–212.

41. Ou H-L, Kim CS, Uszkoreit S, Wickström SA, Schumacher B (2019) Somatic Niche Cells Regulate the CEP-1/p53-Mediated DNA Damage Response in Primordial Germ Cells. Dev Cell 50: 167–183.e8.

42. Zeiser E, Frøkjær-Jensen C, Jorgensen E, Ahringer J (2011) MosSCI and gateway compatible plasmid toolkit for constitutive and inducible expression of transgenes in the C. elegans germline. PLoS One 6: e20082.

43. Sapetschnig A, Sarkies P, Lehrbach NJ, Miska EA (2015) Tertiary siRNAs mediate paramutation in C. elegans. PLoS Genet 11: e1005078.

44. Ge SX, Jung D (2018) ShinyGO: a graphical enrichment tool for ani-mals and plants. bioRxiv 315150.

45. Hakim A, Mor Y, Toker IA, Levine A, Neuhof M, Markovitz Y, Rechavi O (2018) WorMachine: machine learning-based phenotypic analysis tool for worms. BMC Biol 16: 8.

46. Kamath RS, Martinez-Campos M, Zipperlen P, Fraser AG, Ahringer J (2001) Effectiveness of specific RNA-mediated interference through ingested double-stranded RNA in Caenorhabditis elegans. Genome Biol 2: RESEARCH0002.

47. Andrews S (2010) FastQC A Quality Control tool for High Throughput Sequence Data. http://www.bioinformatics.babraham.ac.uk/projects/fastqc/.

48. Langmead B, Salzberg SL (2012) Fast gapped-read alignment with Bowtie 2. Nat Methods 9: 357–359.

49. Anders S, Pyl PT, Huber W (2014) HTSeq A Python framework to work with high-throughput sequencing data. Cold Spring Harbor Labs Journals.

50. Love MI, Huber W, Anders S (2014) Moderated estimation of fold change and dispersion for RNA-seq data with DESeq2. Genome Biol 15: 550.

## Extended Data Reference

43. Zeiser, E., Frøkjær-Jensen, C., Jorgensen, E. & Ahringer, J. MosSCI and gateway compatible plasmid toolkit for constitutive and inducible expression of transgenes in the C. elegans germline. PLoS One 6, e20082 (2011).

44. Sapetschnig, A., Sarkies, P., Lehrbach, N. J. & Miska, E. A. Tertiary siRNAs mediate paramutation in C. elegans. PLoS Genet. 11, e1005078 (2015).

45. Ge, S. X. & Jung, D. ShinyGO: a graphical enrichment tool for ani-mals and plants. bioRxiv 315150 (2018). doi:10.1101/315150

46. Hakim, A. et al. WorMachine: machine learning-based phenotypic analysis tool for worms. BMC Biol. 16, 8 (2018).

47. Kamath, R. S., Martinez-Campos, M., Zipperlen, P., Fraser, A. G. & Ahringer, J. Effectiveness of specific RNA-mediated interference through ingested double-stranded RNA in Caenorhabditis elegans. Genome Biol. 2, RESEARCH0002 (2001).

48. Andrews, S. FastQC A Quality Control tool for High Throughput Sequence Data. http://www.bioinformatics.babraham.ac.uk/projects/fastqc/ (2010).

49. Langmead, B. & Salzberg, S. L. Fast gapped-read alignment with Bowtie 2. Nat. Methods 9, 357–9 (2012).

50. Anders, S., Pyl, P. T. & Huber, W. HTSeq A Python framework to work with high-throughput sequencing data. bioRxiv (Cold Spring Harbor Labs Journals, 2014). doi:10.1101/002824

51. Love, M. I., Huber, W. & Anders, S. Moderated estimation of fold change and dispersion for RNA-seq data with DESeq2. Genome Biol. 15, 550 (2014).

